# A novel infective endocarditis virulence factor related to multiple functions for bacterial survival in blood was discovered in *Streptococcus sanguinis*

**DOI:** 10.1101/2024.07.03.601854

**Authors:** Vysakh Anandan, Liang Bao, Zan Zhu, Jennifer Bradley, Valery-Francine Assi, Henna Chavda, Todd Kitten, Ping Xu

## Abstract

We identified the role of a conserved hypothetical protein (SSA_0451) in *S. sanguinis* that is involved in the virulence of infective endocarditis. An *in vitro* whole blood killing assay and rabbit endocarditis model studies revealed that the SSA_0451 mutant (ΔSSA_0451) was significantly less virulent than the wild-type (SK36) and its complementation mutant (ΔSSA_0451C). The mechanism underlying the SSA_0451 mutant’s reduced virulence in infective endocarditis was evidentially linked to oxidative stress and environmental stress. The genes related to the survival of *S. sanguinis* in an oxidative stress environment were downregulated in ΔSSA_0451, which affected its survival in blood. Our findings suggest that SSA_0451 is a novel IE virulence factor and a new target for drug discovery against IE.

**Author summary:** This study focused on SSA_0451, a conserved hypothetical protein in *S. sanguinis*, to explore its potential role as a virulence factor. Through *in vitro* whole blood killing assays and rabbit IE models, it was found that the SSA_0451 mutant exhibited reduced virulence compared to the wild-type and a complemented mutant. The study linked the mutant’s diminished virulence in IE to heightened susceptibility to oxidative and environmental stresses, supported by downregulation of genes crucial for oxidative stress survival in *S. sanguinis*. These findings identify SSA_0451 as a novel virulence factor in IE and propose it as a promising target for future drug development against this condition.

## Introduction

*Streptococcus sanguinis* is a prevalent oral commensal bacterium and an opportunistic pathogen when the organism gains access to the bloodstream[1–3]. Infective endocarditis (IE) is an infection of the heart’s inner lining (endocardium) or the heart valves. Bacteremia and the presence of endothelial damage are essential factors in the pathogenesis of streptocccal IE[4]. *S. sanguinis* and other oral streptococci are together responsible for an estimated 20% of IE cases worldwide. The initial step in the pathogenesis of IE is the entry of pathogens into the bloodstream. Following entry, the pathogen travels through the blood and reaches the damaged endothelium or valve. Once the bacteria adhere to the damaged endothelium, platelets and fibrin are deposited over the organisms, forming a vegetation. The organisms trapped within the vegetation are protected from phagocytic cells and other host defense mechanisms[4].

*S. sanguinis* undoubtedly posseses virulence factors that contribute to its ability to survive in blood and to cause IE[5]. Given the disparities between the oral cavity, blood, and cardiac vegetation, it is possible that *S. sanguinis* possesses virulence factors that are crucial for IE pathogenesis but not for oral colonization[6]. In *S. sanguinis*, only a few genes necessary for endocarditis virulence have been identified to date[7–11]. Identifying new potential virulence factors contributing to *S. sanguinis* survival in the bloodstream could aid in developing better treatments for controlling IE. Hence, the current study was designed to identify one or more novel IE virulence factor in *S. sanguinis*. The SSA_0451 and SSA_0451 complemented mutant (ΔSSA_0451C) were prepared by PCR-based gene knockout method. RNA-seq was used in this investigation to evaluate the function of this protein. The *in vitro* whole blood killing assay and rabbit IE model studies were used to evaluate its potential as a virulence factor of infective endocarditis. This study also aimed to determine how this novel virulence factor allows *S. sanguinis* to persist in the bloodstream and induce infective endocarditis.

## Results

### Identification of a novel IE virulence factor in a whole-blood-killing assay

In our laboratory, we previously prepared a single gene deletion mutant library of 2046 genes of *S. sanguinis.* We individually cultured each single gene deletion mutant and pooled them based on their equalized OD_600_.The prepared library pool was used for the blood survival experiment (fresh rabbit blood). Using a system biology approach, we identified several mutant with less blood survival potential during these screening study. Based on our preliminary screening data, the ΔSSA_0451 mutant shows less blood survival in genome-wide *in vitro* blood survival screening experiments. Based on this, we designed the current study to explore the potential of SSA_0451 as an IE virulence factor. The ΔSSA_0451 mutant is deleted for a gene annotated as encoding a conserved uncharacterized protein with 129 amino acids. We used the whole blood killing assay to assess the survival capability of the ΔSSA_0451 mutant in blood.

To directly compare the blood killing of ΔSSA_0451 with the wild-type strain SK36, we fluorescently labeled the bacterial cells of SK36, ΔSSA_0451, and ΔSSA_0451C (a complemented mutant) with mCherry (red). These fluorescently labeled strains were used for the flow cytometry analysis to distinguish bacterial cells from the red/white blood cells in the killing assay. Flow cytometry-based analysis revealed a significant reduction of ΔSSA_0451 counts compared to SK36 and its complemented mutant, ΔSSA_0451C (Fig 1; S1 Fig 1). A CFU-based blood-killing assay further validated this finding. Significantly fewer ΔSSA_0451 cells were recovered compared to the complemented mutant (S1 Fig 2). Both results demonstrated that the SSA_0451 gene is essential for *S. sanguinis* survival in blood.

**Fig 1.**
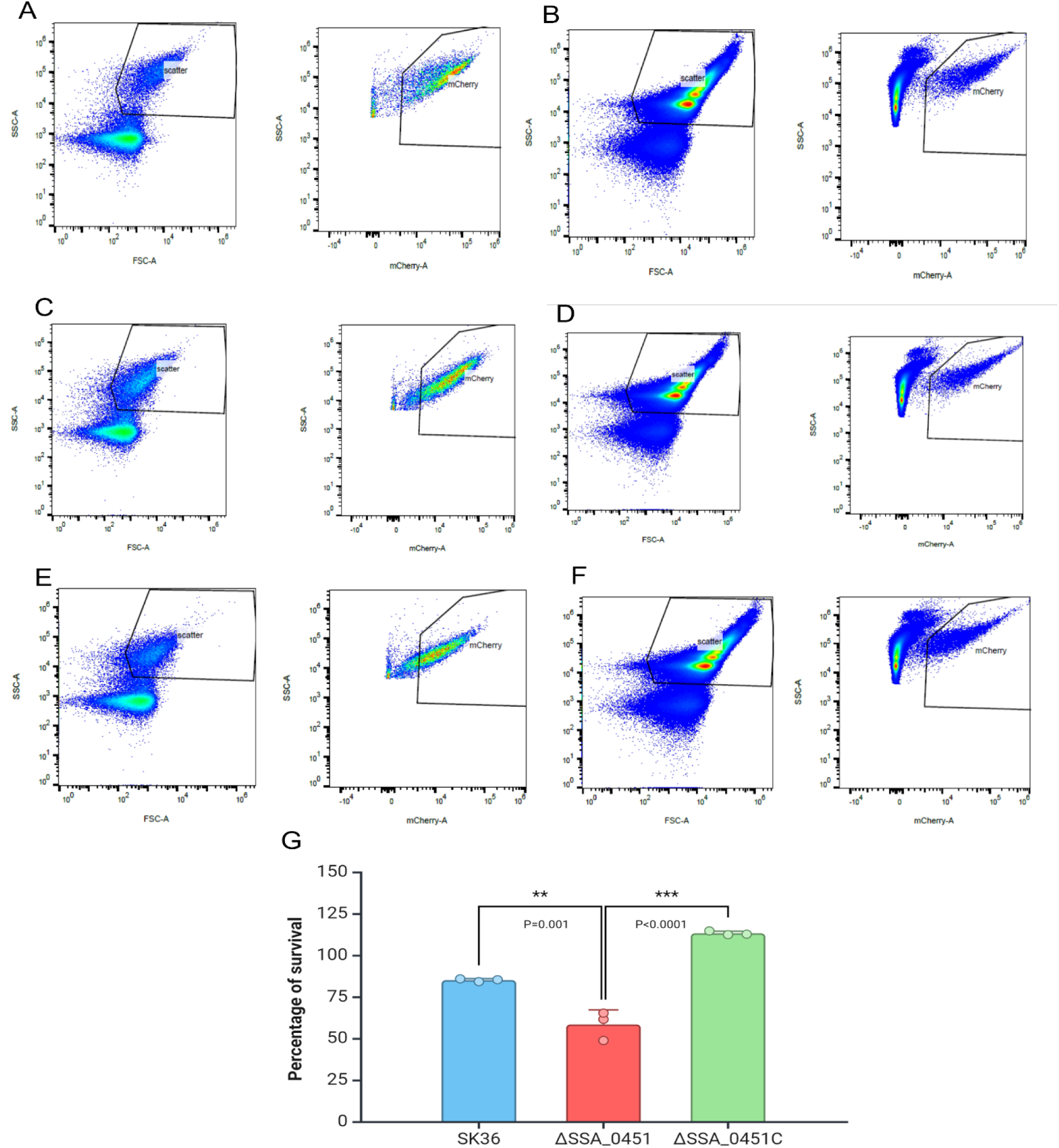
Flow cytometry-based analysis showing the reduced blood survival potential of ΔSSA_0451. SK36, ΔSSA_0451, and ΔSSA_0451C were labeled with mCherry (red). The survival percentage of SK36, ΔSSA_0451, and ΔSSA_0451C were calculated based on the detection of live bacterial cells based on red channel gating. The survival percentage was shown based on before and after treatment bacterial count. ΔSSA_0451 showed reduced blood survival after 30 minutes. A: SK36 T0; B: SK36 T30; C: ΔSSA_0451 T0; D: ΔSSA_0451 T30; E: ΔSSA_0451C T0; F: ΔSSA_0451C T30; G: Percentage of survival of SK36, ΔSSA_0451 and ΔSSA_0451C. Values are ± SD, n=3.

### ΔSSA_0451 shown to be highly sensitive to blood in a competitive index assay

After demonstrating the reduced blood survival of the SSA_0451 mutant, we evaluated its survival potential in an *in vitro* blood-killing assay model using flow cytometry. For this purpose, the wild-type SK36 was labeled with mTFP1 (green) plasmid, while ΔSSA_0451 and ΔSSA_0451C strains were labeled with mCherry (red) plasmid to produce fluorescently labeled strains. The green fluorescence-labeled strain SK36 was mixed with a red fluorescence-labeled strain, either ΔSSA_0451 or ΔSSA_0451C, for a cytometry-based CI assay (Fig 2A-E). The flow cytometry-based CI of ΔSSA_0451 with SK36 was 0.85±0.02, and ΔSSA_0451C with SK36 was 1.04±0.008 respectively, which indicated the virulence attenuation of ΔSSA_0451 (CI <1 indicated the virulence attenuation).

**Fig 2.**
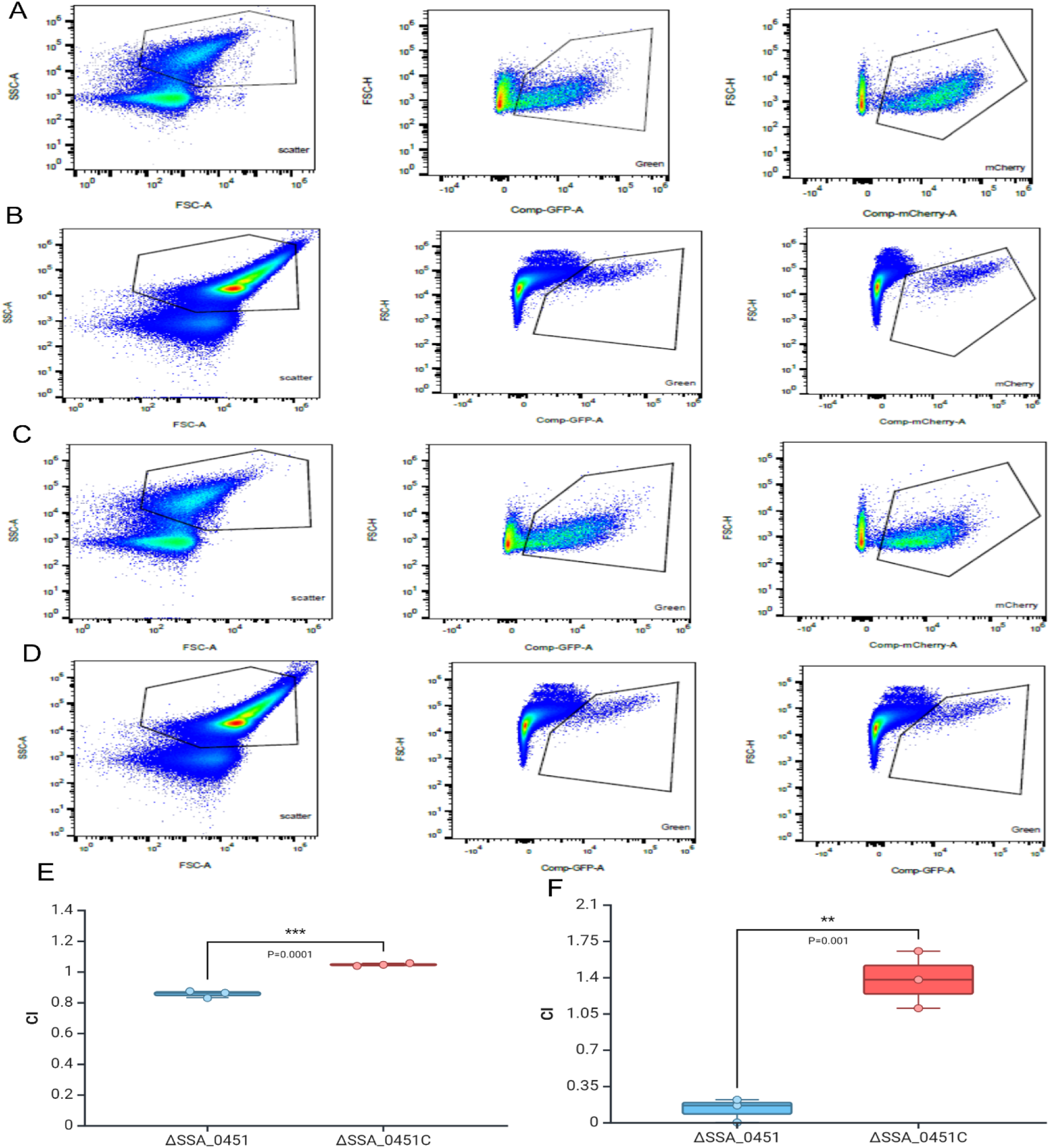
ΔSSA_0451 shows sensitivity to blood in a competitive index assay and virulence attenuation in the rabbit IE model. SK36 was labeled with mTFP1 (green) and mixed with ΔSSA_0451 or ΔSSA_0451C cells (both were labeled with mCherry, red). The CI of ΔSSA_0451 and ΔSSA_0451C were determined in comparison with SK36 based on the live bacteria in red and green channel gating. A:ΔSSA_0451 T0; B: ΔSSA_0451 T30; C: ΔSSA_0451C T0; D ΔSSA_0451C T30; E: CI of ΔSSA_0451 and ΔSSA_0451C from 30 min blood treatment. F: CI of ΔSSA_0451 and ΔSSA_0451C from rabbit endocarditis model experiment. Values are ± SD, n=3. The CI of <1 indicated the virulence attenuation.

The virulence attenuation of ΔSSA_0451 was evaluated using the rabbit IE model. Three different antibiotic markers were introduced into SK36, ΔSSA_0451, and ΔSSA_0451C. An equal number of cells of the three strains were then pooled and injected into catheterized rabbits. The data indicated that the CI for ΔSSA_0451 was 0.12±0.02 (Fig 2F), and for ΔSSA_0451C, it was 1.37±0.008. The CI of ΔSSA_0451 shows a significant difference compared to CI of ΔSSA_0451C (P=0.001). This CI value of SSA_0451 was higher than our previously reported IE virulence factor gene SSA_0260 (one vital virulence factor with CI value of 2.9 x 10^-4^). Additionally, the CI value of SSA_0451 was lower than our previously reported IE virulence factor IspA (CI=0.58) and Igt (CI=0.34)[12]. However, the SSA_0451 mutation had a negative effect through its virulence attenuation (CI<1). Both *in vitro* and *in vivo* approaches demonstrated the function of SSA_0451 as an endocarditis virulence factor.

### RNA Seq analysis indicated the gene expressions of metabolism, ABC transporters, and antioxidant defense system were changed in ΔSSA_0451

Subsequently, we needed to determine how SSA_0451 functions as a virulence factor. To investigate the role of the protein in *S. sanguinis*, we performed an RNAseq analysis of ΔSSA_0451. Global gene expression of ΔSSA_0451 was compared to that of SK36 (S1 Fig 3). A total of 196 genes were downregulated in ΔSSA_0451 with a log_2_ fold change ≤ −1.5, and a total of 75 genes were upregulated with a log_2_ fold change ≥ 1.5 (P≤ 0.05). A pathway enrichment analysis was performed with the KEGG and GO databases. The pathways enriched include fatty acid metabolism, galactose metabolism, and biosynthesis of cofactors (S1 Fig 4). The STRING network analysis revealed an enrichment of pathways linked to fatty acid metabolism (Fig 5IA-C), lactose metabolism (Fig 5IIA-C), PTS-system-mannose specific (S1 Table 1), thiol peroxidase-associated genes (Fig 3B), and ABC-type Mn/Zn transporters (Fig 4). The pathways related to cellular response to stimulus, competence and transformation-related genes were upregulated (Fig 5IIIA-C) with log2 fold change ≥ 1.5.

**Fig 3.**
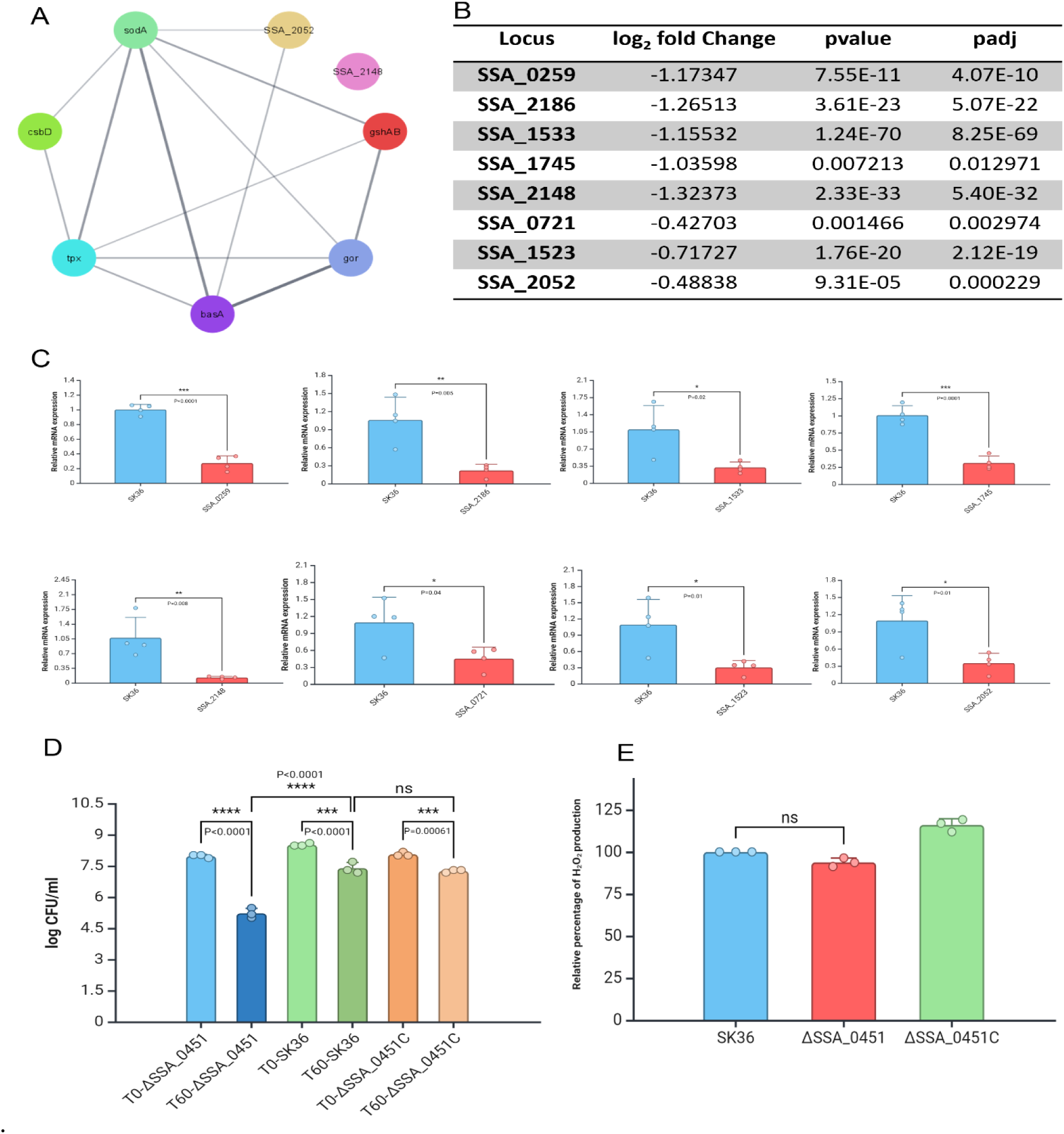
The SSA_0451 gene is necessary for *S. sanguinis* to survive under oxidative stress. A: String network of genes involved in thiol-specific peroxidase mediated detoxifying process and antioxidant-related genes in SK36. B: RNA-seq data showing the deletion of the SSA_0451 gene downregulates the genes involved in the peroxidase detoxifying process and antioxidant protection mechanism. C: RNA-seq data were validated with qRT-PCR. D: Deleting the SSA_0451 gene reduced the survival of ΔSSA_0451 in an H_2_O_2_-mediated stress environment. E: Deleting the SSA_0451 doesn’t impact the H_2_O_2_ production capability of *S. sanguinis*. The antioxidant activity-related genes and thiol peroxidase-related genes help the *S. sanguinis* to thrive in an oxidative stress environment. Downregulation of these genes made ΔSSA_0451 susceptible to oxidative stress.

Fatty acid biosynthesis is an essential activity in most bacteria, which must be regulated by precursor availability and cell division requirements. The carboxylation of acetyl-CoA into malonyl-CoA is the initial step in fatty acid biosynthesis. The CoA is then replaced with an acyl carrier protein (ACP). Malonyl-ACP acts as a universal substrate for fatty acid elongation by sequentially adding acetyl-moieties. The ACCase complex consists of four subunits grouped into two subcomplexes: AccB-AccC and AccA-AccD. The initial stage in the reaction is the carboxylation of biotin by biotin carboxylase (AccC). The biotin cofactor binds to the biotin-carboxyl carrier protein (AccB) [13]. Finally, ACCase’s carboxyltransferase domain (AccAD) transfers the carboxyl group to acetyl-CoA [14]. All ACCase subunits are important for membrane biosynthesis and have been considered potential therapeutic targets[15]. In our RNA-SEQ data, the deletion of SSA_0451 leads to the downregulation of all subunits of ACCase, such as SSA_1930 (accA), SSA_1931 (accD), SSA_1932 (accC), SSA_1934 (accB). The other genes related to fatty acid biosynthesis include SSA_1933 (FabZ), SSA_1935 (FabF), SSA_1936 (FabG), SSA_1937 (FabD), SSA_1938 (FabK), SSA_1939 (acpP-Acyl carrier protein), and SSA_1940 (FabH) were downregulated in ΔSSA_0451 (Fig 5IA-C). These observations confirmed that the deletion of the SSA_0451 gene affects the fatty acid biosynthesis pathway of *S. sanguinis*.

The phosphoenolpyruvate (PEP):carbohydrate phosphotransferase system (PTS) plays a crucial role in the regulation of central carbon metabolism in *S. sanguinis*, which contributes to the ability of this commensal bacterium to tolerate stress and compete against *S. mutans*[16]. According to our RNA-Seq data, the SSA_0451 gene deletion results in the downregulation of genes linked to a cluster of genes of the PTS system’s mannose-specific IIC component, including SSA_0219, SSA_0220, SSA_0221, SSA_0222, SSA_2023, and SSA_0224 (Extended data table 1).The lactose metabolism-related genes like SSA_1692, SSA_1693, SSA_1694, SSA_1695, SSA_1696, SSA_1697, SSA_1698, SSA_1699, SSA_1700 were downregulted in ΔSSA_0451. The gene SSA_1701 (lactose repressor LacR) functions as a transcriptional repressor of sugar metabolism in *S. sanguinis* and regulates the catabolism of multiple carbohydrates. The deletion of SSA_0451 downregulated the expression of LacR by −0.68929 log2 fold. Comparing the growth and carbohydrate transport in the *lacR* mutant of SK36 to other *lacR* mutants, Zeng et al., observed an exceptionally severe defect in PTS activity compared to similarly constructed *lacR* mutants[17]. Moreover, the glucose-PTS gene (SSA_1918, *manL*) that encodes the A and B domains of a PTS permease enzyme II was downregulated (−0.89121 log2 fold) in the SSA_0451 mutant. This downregulation of *manL* could reduce the capacity of *S. sanguinis* to utilize several carbohydrates commonly present in the human oral cavity. These data revealed that the SSA_0451 is essential for *S. sanguinis* to survive in nutrient-limited environments. In *Streptococcus agalactiae* and *Streptococcus pneumoniae,* similar behavior was reported in these families of proteins[18,19]. Two genes that encode putative complement system proteases are present in the genome of *S. sanguinis* SK36: *pepO* (encoding endopeptidase O) and *cppA* (annotated as C3-complement degrading protease; CppA)[20]. PepO homologs in *S. pneumoniae*, *Streptococcus pyogenes*, and *Streptococcus mutans* are responsible for complement evasion and invasion into host cells [21–24]. Research on *S. pneumoniae* and *S. pyogenes* further points to potential roles for CppA in bacterial virulence [25–27].. According to a study conducted by Lívia Alves et al., *S. sanguinis* expresses complement evasion proteins like PepO and CppA in a strain-specific manner that are necessary for various functions related to cardiovascular virulence[28]. The SSA_0331 gene codes for CppA, while the SSA_0263 gene codes for PepO proteins in *S. sanguinis*. The expressions of two complement evasion proteins, PepO (−1.18 log2 fold) and CppA (−0.53 log2 fold), were downregulated by the deletion SSA_0451 gene. The gene downregulations could be one of the possible explanations for SSA_0451’s decreased virulence in the endocarditis model.

### The SSA_0451 gene is necessary for *S. sanguinis* to survive under oxidative stress

Thiol-specific peroxidase catalyzes the reduction of hydrogen peroxide and organic hydroperoxides to water and alcohol and plays a role in cell protection against oxidative stress by detoxifying peroxides. In *S. sanguinis*, the deletion of the SSA_0451 gene downregulates SSA_0259, SSA_2186, SSA_1533, SSA_1745, SSA_2148, SSA_0721, SSA_1523, SSA_2052 genes which are involved in the peroxidase detoxifying process (Fig 3A). Moreover, the enzyme that protects *S. sanguinis* from ROS-mediated stress was downregulated upon deletion of the SSA_0451 gene (Fig 3B). Data from our qRT-PCR study supports these findings (Fig 3C). The downregulated antioxidant enzyme-related genes in ΔSSA_0451 provide a better environment for the immune cells to clear the bacterium from the blood. Subsequently, we evaluated the H_2_O_2_ production and H_2_O_2_ survival potential of SSA_0451 to test this hypothesis. Sumioka et al., reported that *S. sanguinis* induces neutrophil cell death by H_2_O_2_ production[29]. In our H_2_O_2_ production assay, the deletion of SSA_0451 doesn’t impact the H_2_O_2_ production capability of *S. sanguinis* (Fig 3E). Interestingly, the H_2_O_2_ survival was reduced in ΔSSA_0451 compared to ΔSSA_0451C and SK36 (Fig 3D). These data also strengthen our concept that the reduced blood survival might be due to ROS-mediated killing.

The lipoprotein SsaB, in conjunction with the SsaACB transporter complex, plays a crucial role in the virulence of *S*. *sanguinis* for endocarditis[12]. SsaB, belonging to the LraI family of conserved metal transporters, has been identified as critical for *S. sanguinis* virulence for endocarditis[6,12]. Besides, manganese (Mn) is an essential nutrient for *S. sanguinis*, and its acquisition through the SsaB protein enhances the bacterium’s ability to resist oxidative stress and survive within the host. Studies have shown that intracellular Mn accumulation mediated by SsaB promotes virulence and O_2_ tolerance of *S. sanguinis*[30]. The SsaACB manganese transporter complex has been established as critical for redox maintenance in *S. sanguinis*[30]. In addition, our RNA-seq data showed that the ABC-type Mn transporters (SSA_0260, SSA_0261, SSA_0262) were substantially downregulated during SSA_0451 deletion (Fig 4). The SsaACB transporter complex downregulation in ΔSSA_0451 could be the another reason behind its reduced virulence in IE.

**Fig 4.**
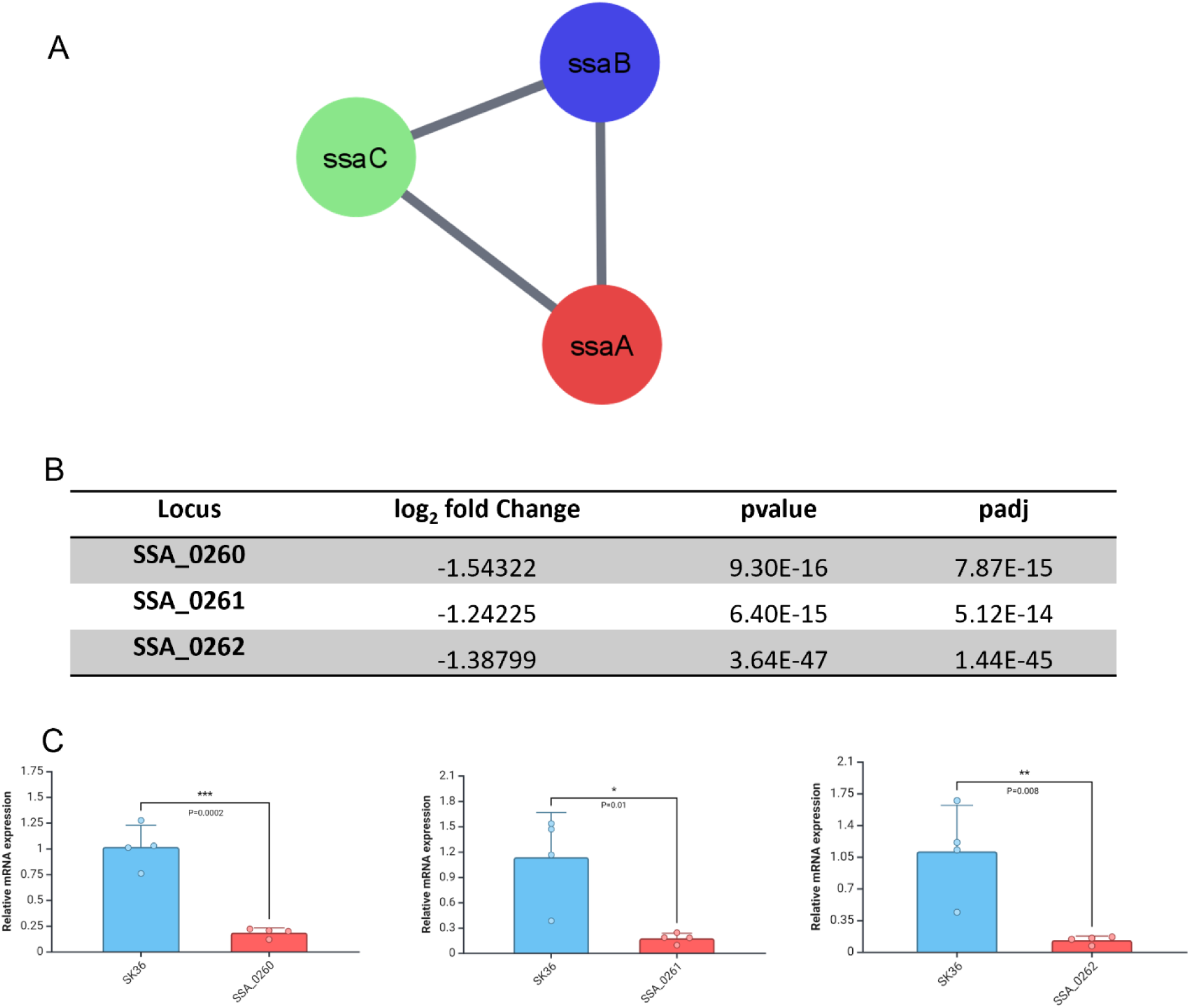
ABC-type Mn/Zn transporters were downregulated during SSA_0451 gene deletion. A: Mn/Zn transporters-related gene network. B: RNA-seq data shows the downregulation of ABC-type Mn/Zn transporters-related genes during the SSA_0451 gene deletion. C: qRT-PCR validation of RNA-seq data shows the downregulation of SSA_0260, SSA_0261, and SSA_0262 genes.

**Fig 5.**
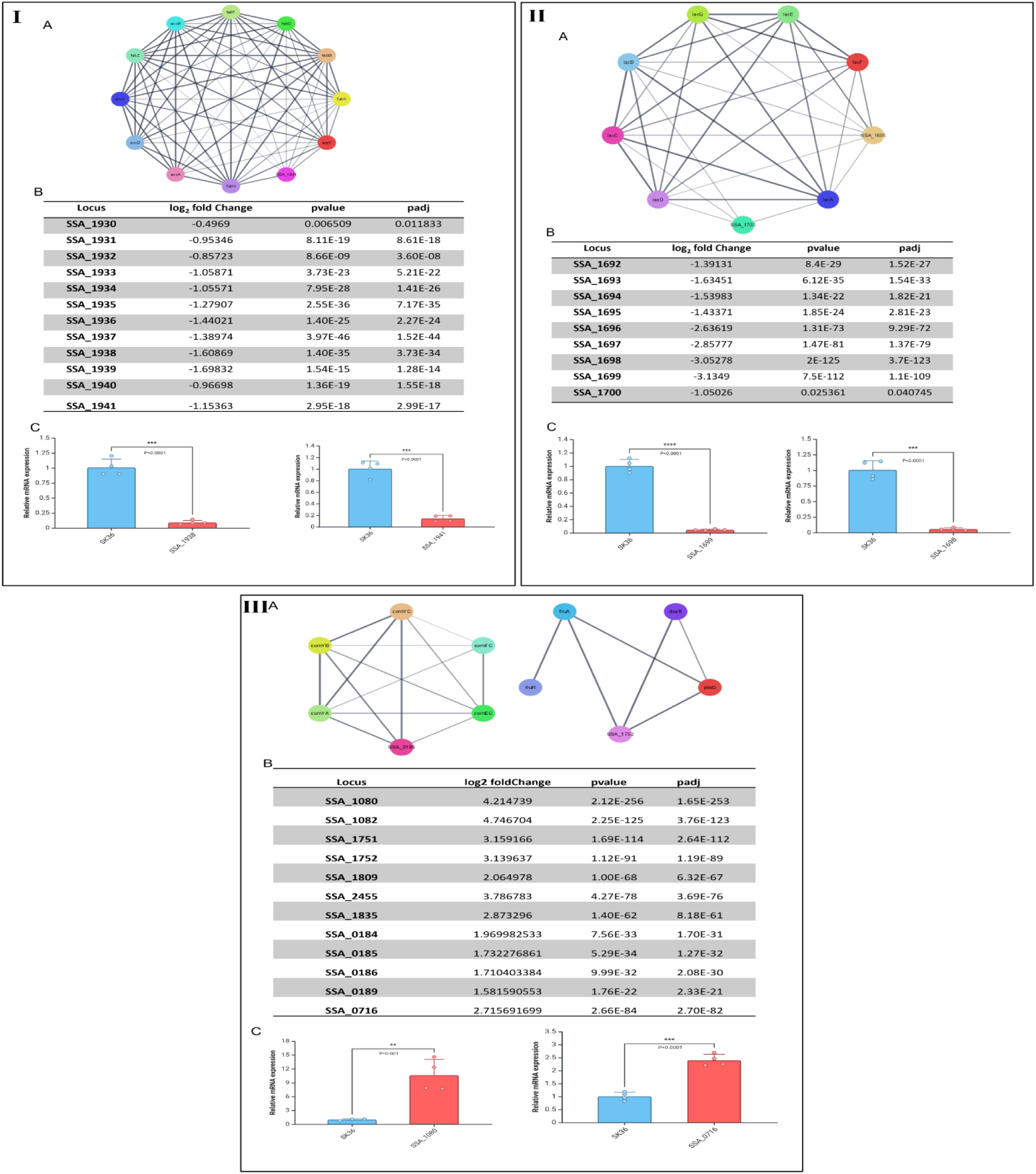
The SSA_0451 gene is necessary for *S. sanguinis* to survive under environmental stress. I-A,B,C: Fatty acid metabolism-related gene network, RNA-seq data, qRT-PCR validation of RNA-seq data respectively. II-A,B,C: Lactose metabolism-related gene network, RNA-seq data, qRT-PCR validation of RNA-seq data respectively. III-A,B,C: competence and fructose metabolism-related genes network, RNA-seq data, qRT-PCR validation of RNA-seq data respectively.

### The SSA_0451 gene is necessary for *S. sanguinis* to survive under environmental stress

To understand SSA_0451’s role in surviving unfavorable environmental conditions, the growth and survival potential of ΔSSA_0451, ΔSSA_0451C, and SK36 were assessed in various environmental conditions, including pH, bile salt, and H_2_O_2_ challenges. The survival rate of ΔSSA_0451 was significantly decreased compared to SK36 in acidic conditions at pH-3 (S1 Fig 5-IA). Additionally, there was no significant growth difference between ΔSSA_0451C and SK36 was observed at pH-3. At pH-7.4, the ΔSSA_0451 and SK36 showed no significant difference in their growth (S1 Fig 5-IB). The alkaline condition favored the growth of ΔSSA_0451C and SK36 but not ΔSSA_0451 growth (pH-9.5) (S1 Fig 5IC). These findings demonstrate that *S. sanguinis* needs SSA_0451 in order to maintain growth in an alkaline environment (pH-9.5). Kelsi L. Anderson et al., reported similar findings on alkaline stress response in *Staphylococcus aureus*[31]. These observations demonstrated that the SSA_0451 protein might belong to the Gls24 or alkaline shock protein-23 family.

The bile salt tolerance potential of ΔSSA_0451, ΔSSA_0451C, and SK36 was analyzed up to 24 hours at 37°C, aerobically. The survival of ΔSSA_0451 in bile salt was significantly reduced compared to SK36 and ΔSSA_0451C (S1 Fig 5II). In *Enterococcus faecalis,* Asp-23/Gls24 family protein was shown to be induced under other stress conditions, such as the presence of bile salt and cadmium chloride, so it was designated as a general stress (Gls) protein[32]. Our complementation and mutation data revealed that SSA_0451 is involved in *S. sanguinis’s* ability to tolerate bile salts and general stress.

## Discussion

The opportunistic bacterium *S. sanguinis*, which survives within the bloodstream and gains access to heart valves and causes IE[33,34]. *S. sanguinis* virulence factors that contribute to the pathogenic process are poorly understood, and many potential virulence genes have not been experimentally examined. We designed the current study to investigate the potential of SSA_0451 as an IE virulence factor. Our detailed investigation found that the conserved uncharacterized protein SSA_0451 was a novel IE virulence factor, and it was essential for *S. sanguinis* to survive in different environmental conditions.

Initially, we investigated the potential of the SSA_0451 gene as a virulence factor of IE. The potential of SSA_0451 as a significant virulence factor in IE was demonstrated by the CI measurements and the *in vitro* blood-killing assays. The *in vivo* findings substantiated the *in vitro* research and demonstrated that SSA_0451 is a crucial virulence factor that aids *S. sanguinis* in surviving in a blood environment. It is the first study to report that SSA_0451 is an IE virulence factor.

Because SSA_0451 is a conserved hypothetical protein, and we need to determine its precise function. A careful protein database search indicated that the SSA_0451gene sequence shows similarities to Asp23protein family(S1 Fig 6). Asp23 is one of the poorly investigated proteins belonging to the Asp23 protein family, also termed DUF322, PF03780, or Gls24 family[35]. Members of the Gls24 family proteins are commonly present in Gram-positive bacteria [36]. This family of proteins was functionally associated with envelope stress in *Staphylococcus aureus* [36]. In *Bacillus subtilis,* these proteins were linked to lipid metabolism [37]. In *Streptococcus agalactiae,* this stress protein was necessary for bacteria to thrive in low pH and nutrient-limited environments [18]. The bile stress response, cell morphology control, and intestinal environment adaptation in *Enterococcus faecium* were controlled by Gls24 stress protein family proteins [32,38] The nutrition-sensing process of *S. pneumoniae* was significantly influenced by this protein family[19]. Our protein database search, RNA-Seq analysis (reduced lipid metabolism genes, nutrition-related, increased stress, and competence-related genes), and environmental stress experiments (evidenced by its reduced survival in pH, bile salt, and H_2_O_2_), all pointed to the conclusion that SSA_0451 is an Asp23 family protein and related to the general stress response of *S. sanguinis.* The discovery of SSA_0451 as an IE virulence factor paves the way for IE drug development as a new pharmacological target. By regulating the SSA_0451 gene, we can control multiple pathways necessary for SK36 survival in blood and adverse environments.

**Fig 6.**
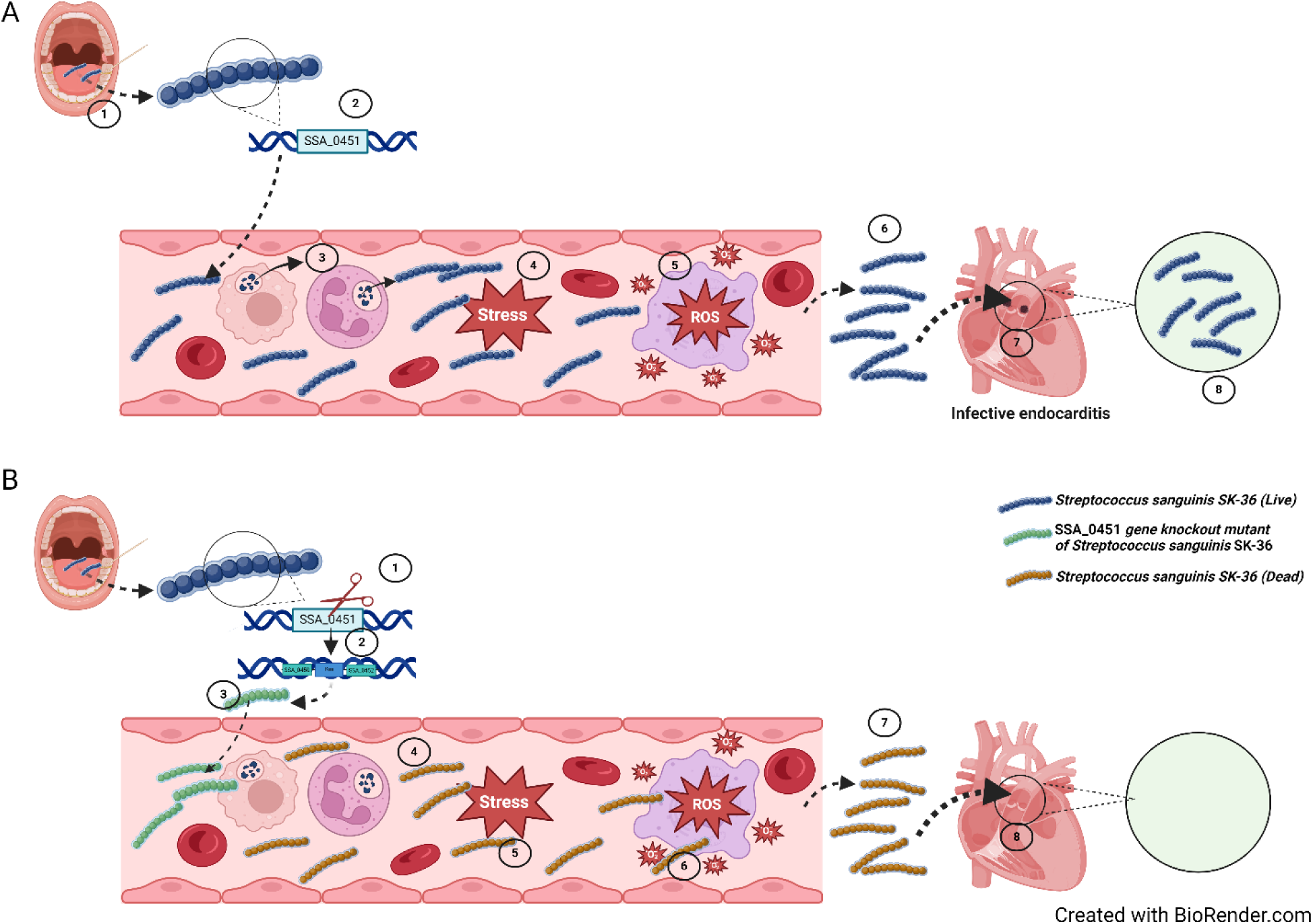
A schematic illustration of the SSA_0451 gene’s role in *S. sanguinis-*mediated infective endocarditis. **A1**: The opportunistic pathogen *S. sanguinis* SK36 gains access to the bloodstream during dental procedures. **A2**: The SSA-0451 gene is one of the hypothetical protein genes present in SK36, and this gene’s function in IE is still unknown. **A3, A4 and A5**: During the SK36 entry into the blood, the ROS generated from immune cells create a stressful environment within the bloodstream. **A6**: The antioxidant system-related genes and thiol peroxidase-related genes help the SK36 to thrive from ROS-mediated stress. **A7**: The blood survived SK36 colonized on the damaged heart valve, causing IE. **B1:** The deletion of the SSA_0451 gene from SK36 was carried out using a PCR-based method. **B2:** The SSA_0451 gene is replaced with the Km gene and made SSA_0451 gene knockout mutant of SK36. **B3:** The ΔSSA_0451 enters the bloodstream. **B4:** Once entered into the bloodstream, the ROS-mediated killing occurs by the action of immune cells. **B5:** The SSA_0451 gene is needed for the SK36 to survive in environmental and ROS-mediated stressful conditions. **B6:** Due to the deletion of the SSA_0451 gene, the ΔSSA_0451 could not survive in the adverse environmental conditions. **B7 and B8:** The ΔSSA_0451 was easily cleared by blood cells, making the ΔSSA_0451 incompetent to cause IE.

Moreover, one direct link to the reduced blood survival of SSA_0451 was revealed from our RNA-Seq data.Lívia Alves et al., reported that PepO and CppA are complement evasion proteins expressed by *S. sanguinis* in a strain-specific manner, which are required for multiple functions associated with cardiovascular virulence[24,28]. The genes related to complement evasion, such as SSA_0331 (CppA) and SSA_0263 (PepO) in *S. sanguinis,* were downregulated by deletion of the SSA_0451 gene. Additionally, deleting the SSA_0451 gene downregulates genes involved in the peroxidase detoxifying process and upregulates the genes responsible for H_2_O_2_ production. These data suggest that the reduced blood survival might be due to ROS-mediated killing. Our H_2_O_2_ survival study supports this hypothesis, demonstrating that ΔSSA_0451 has lower survival rates than SK36.

After analyzing the combined data, we developed a model of how the SSA_0451 gene may contribute to *S. sanguinis-*mediated infective endocarditis (Fig 6). Once the opportunistic pathogen *S. sanguinis* SK36 gains access to the bloodstream, it utilizes different kinds of virulence factors and various mechanisms to escape from blood-mediated killing. The ROS generated from immune cells creates a stressful environment within the bloodstream. The antioxidant system-related genes and thiol peroxidase-related genes help SK36 to survive ROS-mediated stress, allowing bacteria to reach the damaged heart valve and colonize it, resulting in IE. Deleting the SSA_0451 gene from SK36 downregulated the antioxidant system-related genes and thiol peroxidase-related genes, making ΔSSA_0451 unable to survive in the adverse environmental conditions in blood. Thus, ΔSSA_0451 was easily cleared by blood cells and was thus incompetent to cause IE.

## Methods

### Collection of Blood sample

For blood collection, pathogen-free New Zealand white rabbits weighing 3–4 kg were purchased from Charles River and allowed to acclimate to the vivarium before blood collection. In keeping with guidelines, we collected ≤ 10% blood volume per draw and waited ≥ 2 weeks before drawing blood again from the same animal. All animal experiments were performed in accordance with the guidelines of the U.S. Office of Laboratory Animal Welfare and the U.S. Department of Agriculture. All procedures for animal experiments were approved by the Virginia Commonwealth University Institutional Animal Care and Use Committee (Protocol AM10030).

### Mutant and complementation mutant construction

A human oral isolate strain of *S. sanguinis* (*S. sanguinis* SK36) was obtained from Mogens Killian (University of Aarhus, Denmark)[39]. The strains and plasmids used for the study are included in S1 table 2. The single gene deletion of SSA_0451 in *S. sanguinis* SK36 previously constructed in our laboratory was used for this study[40]. For complementation of ΔSSA_0451, a similar [40] PCR-based method was employed [41]. Briefly, the region upstream of SSA_0451, the region downstream of the same gene, and the *aad9* gene encoding resistance to spectinomycin were independently amplified using appropriate primer sets (S1 table 3). The SSA_0451CF1/SSA_0451CR1 was used for the amplification of 1-kb sequence upstream plus the coding sequence of SSA_0451. The SpecF2/SpecR2 (For Spectinomycin (Sc) selection) or EmF2/EmR2 (For Erythromycin (Em) selection) was used for the amplification of the erythromycin resistance cassette (pVA838) or spectinomycin cassette (pDR111) [42]. SSA_0451CF3/SSA_0451CR3 was used to amplify the 1-kb sequence downstream of SSA_0451. Overlapping PCR generated the final recombinant PCR product that contained these three fragments and was then further introduced into ΔSSA_0451 to replace the kanamycin resistance (Km) cassette with the SSA_0451 ORF and the erythromycin or spectinomycin resistance cassette. An erythromycin or spectinomycin-resistant and kanamycin-sensitive transformant was selected and confirmed by PCR and sequence analysis.

### Whole Blood killing assay of ΔSSA_0451

A flow cytometry method was used to analyze bacterial survival for the whole-blood killing assay. The ΔSSA_0451(Km resistant), ΔSSA_0451C (Sc resistant), and WT (SK36) were labeled with mCherry plasmid (containing Em gene as selection marker). The red fluorescent protein mCherry was expressed from the plasmids pVMcherry (S1 Table 2) [43]. The transformed colonies of red fluorescent-producing ΔSSA_0451, ΔSSA_0451C, and SK36 were identified from the antibiotic selection plate. The red fluorescence of these strains was further confirmed with epi-fluorescent microscopy and flow cytometry. The Lancefield bactericidal assay was performed with minor modifications[44]. Briefly, the freshly collected rabbit blood was mixed with 1/10 volume of mCherry labeled SK36, mCherry labeled ΔSSA_0451, and mCherry labeled ΔSSA_0451C (mid-log phase) individually in different tubes. After incubation at 30 minutes, the mixtures were serially diluted with cold, sterile water to lyse the blood cells. 1x RBC lysis buffer (Invitrogen) was used to remove the RBC. The data acquisition was performed using a 5-laser Cytek Aurora Spectral Analyzer with a SSC threshold. Data was analyzed using FlowJo, version 10.8.0.

### Competitive index of ΔSSA_0451 in blood

Flow cytometry method was used to analyze the CI of ΔSSA_0451 in blood. For the flow cytometry experiment, we labeled the SK36 with an mTFP1 plasmid (which produced a green fluorescent protein and contained the Em gene as a selection marker). The equal volume OD_600_ (0.5 OD) of mid-log phase culture of red fluorescent protein producing ΔSSA_0451 mixed with green fluorescent protein producing SK36. The mixture culture was treated with rabbit blood for 30 minutes; the mixtures were then serially diluted with cold sterile water to lyse the blood cells. The samples before and after treatment were analyzed using the Cytek Aurora CS flow cytometry instrument (red and green gating). The data were further analyzed using the flowjo software. The CI was calculated as the ratio of mutant/JFP36 in time zero divided by the mutant/JFP36 ratio after 30 min.

### Rabbit infective endocarditis model

The rabbit model of infective endocarditis was used for testing the virulence of ΔSSA_0451[45]. The rabbits were surgically catheterized (19-gauge catheter-BD Bioscience) through the right internal carotid artery past the aortic valve to cause minor damage. The catheter remained in the artery for the entire experiment. Two days following catheterization, the mid-log culture of SSA_0451(Km), SSA_0451C (Em), and Sc-resistant wild type (JFP56) strains were co-inoculated via peripheral ear vein. After that, the inoculum was diluted and plated on BHI agar with appropriate antibiotics for bacterial counts. At 20 h post-inoculation, the vegetation was collected from rabbit hearts after the verification of catheter placement. Vegetations were homogenized with PBS and plated on BHI agar with appropriate antibiotics for bacterial counts. The CI value for each rabbit was calculated as the ratio of mutant/JFP36 in the vegetation homogenate divided by the mutant/JFP36 ratio in the inoculum. All animal experiments were performed in accordance with the guidelines of the U.S. Office of Laboratory Animal Welfare and the U.S. Department of Agriculture. All procedures for animal experiments were approved by the Virginia Commonwealth University Institutional Animal Care and Use Committee (Protocol AM10030).

### RNA Seq of ΔSSA_0451

ΔSSA_0451 and *S. sanguinis* SK36 were cultured in BHI medium overnight, diluted into fresh BHI medium, and grown for another 4 hours under microaerobic conditions at 37 °C. Samples were collected and treated with RNA protect bacteria reagent (Qiagen, Valencia, CA) for 5 min to stabilize RNA and stored at −80 °C. Cells were lysed by mechanical disruption using FastPrep lysing matrix B (Qbiogene, Irvine, CA). Total RNA was treated with DNase I (Qiagen) and prepared using RNA easy mini kits (Qiagen) according to the manufacturer’s instructions. Total RNA quantity and integrity were determined using a Bioanalyzer (Agilent). All samples passed quality control assessment with RNA Integrity Numbers (RIN) above 8. rRNA depletion was carried out from 100ng of total RNA with NEBNext bacterial rRNA depletion kit (NEB#e7850). NEBNext Ultra II Directional RNA Library Prep Kit for Illumina (New England BioLabs) was used for the following RNA-seq library preparation according to the manufacturer’s protocol. The Bioanalyzer and Tapestation data were used to determine the size distribution (300 bp) and concentration of libraries. Library sequencing was performed by the Nucleic Acids Research Facilities at Virginia Commonwealth University using Illumina HiSeq2000. Reads obtained from RNA-seq were aligned against the *S. sanguinis* SK36 genome using EDGE-pro [46]. Gene sequence reads for *S. sanguinis* SK36 were compared to the ΔSSA_0451 sample using DESeq2 [47] in R (version 4.0.5) to determine log2 fold changes and adjusted p-values. Principal component analysis was completed using R (version 4.0.5) and RStudio (version 13.959)[48].

### Validation of RNA-Seq data by qRT-PCR

qRT-PCR was performed as described previously [41]. First-strand cDNA synthesis was performed in a 20 μl system containing 100 ng Total RNA, 0.2 μl Random Primer (3.0 μg/μl), 1.0 μl dNTP (10 mM each dNTP), 1.0 μl 100 mM DTT, 1.0 μl RNase OUT (40 U, Invitrogen) and 1.0 μl SuperScript III reverse transcriptase (200 U, Invitrogen), 4.0 μl first strand buffer and RNase free water to a 20 μl volume. First-strand cDNA from each reaction was subjected to 10-fold dilution and used in subsequent qRT-PCR. The qRT-PCR was prepared in reactions containing 10 μl 2X SYBR Green PCR Master Mix (Applied Biosystems, Foster City, CA), 0.5 μl each PCR primer (20 μM) (S1 Table 4), 1 μl diluted first strand cDNA and distilled water to a 20 μl volume. The reaction was performed on an ABI 7500 fast real-time PCR system. The housekeeping gene *gyrA* was used as a normalization control. The 2^-ΔΔCt^ method was employed to calculate the relative expression levels of target genes.

### Bacterial growth and survival in different environments

The growth and survival of ΔSSA_0451, ΔSSA_0451C, and SK36 at different pH, Bile salt, and H_2_O_2_ were analyzed using different methods. The growth of ΔSSA_0451, ΔSSA_0451C, and SK36 at different pH were measured aerobically in 96-well plates at 37°C. The bile salt tolerance was measured by adding bile salt at 1mg/ml concentration to BHI media containing ΔSSA_0451, ΔSSA_0451C, and SK36 in 96-well plates. For both experiments, the growths were monitored at 600 nm (up to 24 hrs) with a Synergy H1 Hybrid Reader (BioTek, USA). Three replicates were examined to calculate the means and standard deviations.

For the H_2_O_2_ sensitivity assay, the exponential growth phase cells of SK36, ΔSSA_0451, and ΔSSA_0451C cultured micro-aerobically in BHI broth were collected and washed with sterile PBS for stress treatment. The cell suspension was treated with 20mM H_2_O_2_ (Invitrogen) at 37°C for 60 min and then serially diluted in PBS. The serial dilutions were plated on BHI agar, and the colonies were counted after a 2-day incubation. The bacterial survival was expressed by the percentage of CFU in the treated versus untreated cells.

The H_2_O_2_ production of ΔSSA_0451, ΔSSA_0451C, and SK36 was quantified using an Amplex^TM^ Red Hydrogen Peroxide/Peroxidase kit (Invitrogen, CA). Final values are shown relative to that of the wild-type strain, SK36.

## Contributions

VA. and P.X. conceived and designed this study. VA. carried out all of the experiments with the assistance of L.B. and J.B, T.K, Z.Z, V.F.A, H.C., V.A. and P.X. analyzed the data and wrote this manuscript. All authors reviewed and discussed the manuscript.

## Acknowledgments

We thank Valery-Francine Assi, Jasmine Benbei, and Nicai Zollar for assistance with the animal surgeries, and phlebotomy. Flow cytometry services in support of the research project were provided by the VCU Massey Cancer Center Flow Cytometry Shared Resource supported, in part, with funding from NIH-NCI Cancer Center Support Grant P30 CA016059. The sequence data included in this study was generated at the Genomics Core facility at Virginia Commonwealth University. RNA-Seq analysis was performed on servers provided by the Center for High-Performance Computing at VCU. This work was supported by National Institutes of Health grants NIH-NIDCR R01DE030121 (PX and TK). The funders had no role in study design, data collection, and interpretation, or the decision to submit the work for publication.

## Declaration of interests

The authors declare no competing interests.

---

**S1 Fig 1.**
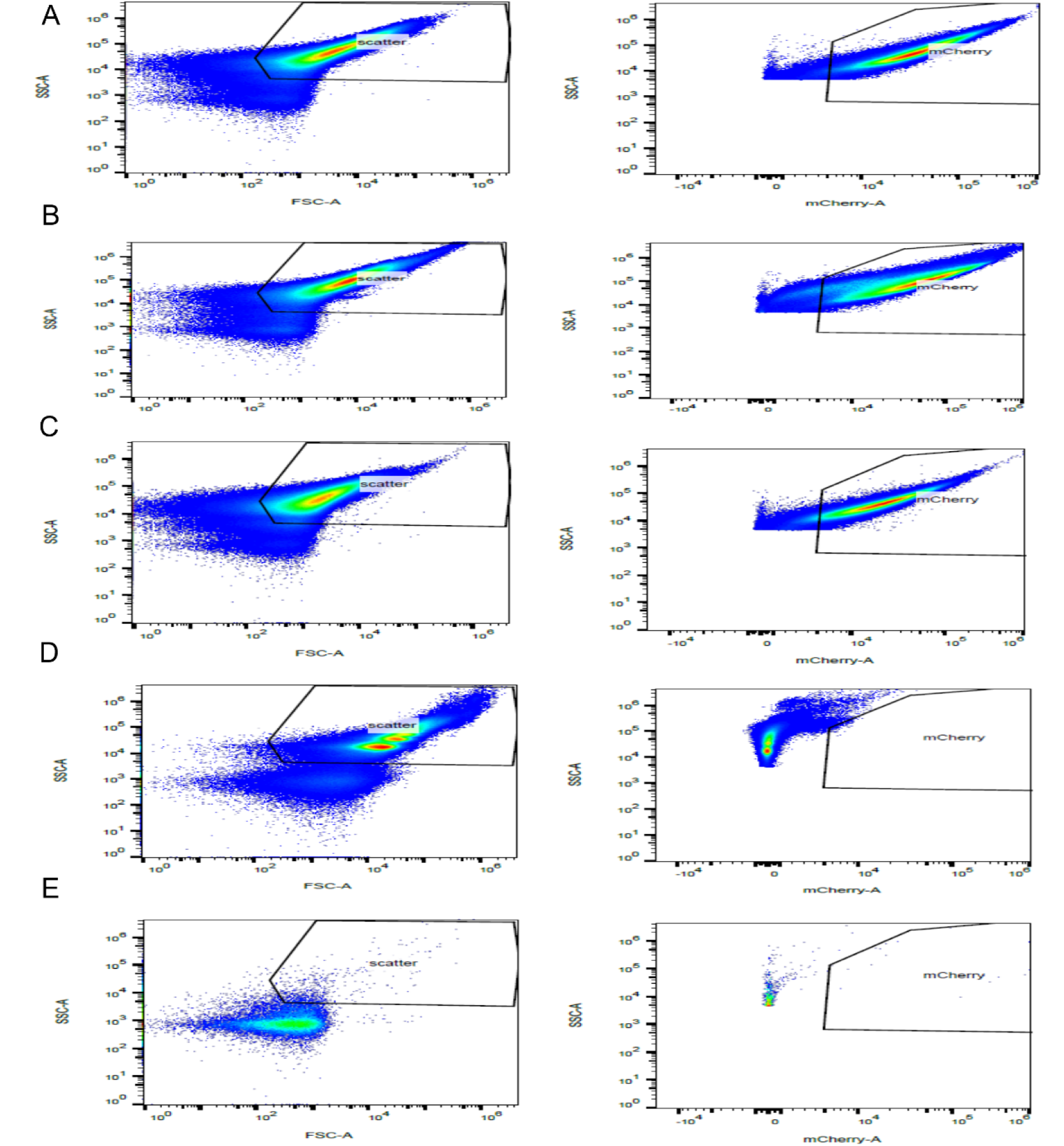
Gate setting on Flow cytometry based on different controls. A: Red colored ΔSSA_0451 appeared in mCherry gate; B: Red colored SK36 appeared in mCherry gate; C: Red colored ΔSSA_0451C appeared in mCherry gate; D: Blood only control not appear in mCherry gates; E: Sterile water control also not appeared in mCherry gates.

**S1 Fig 2.**
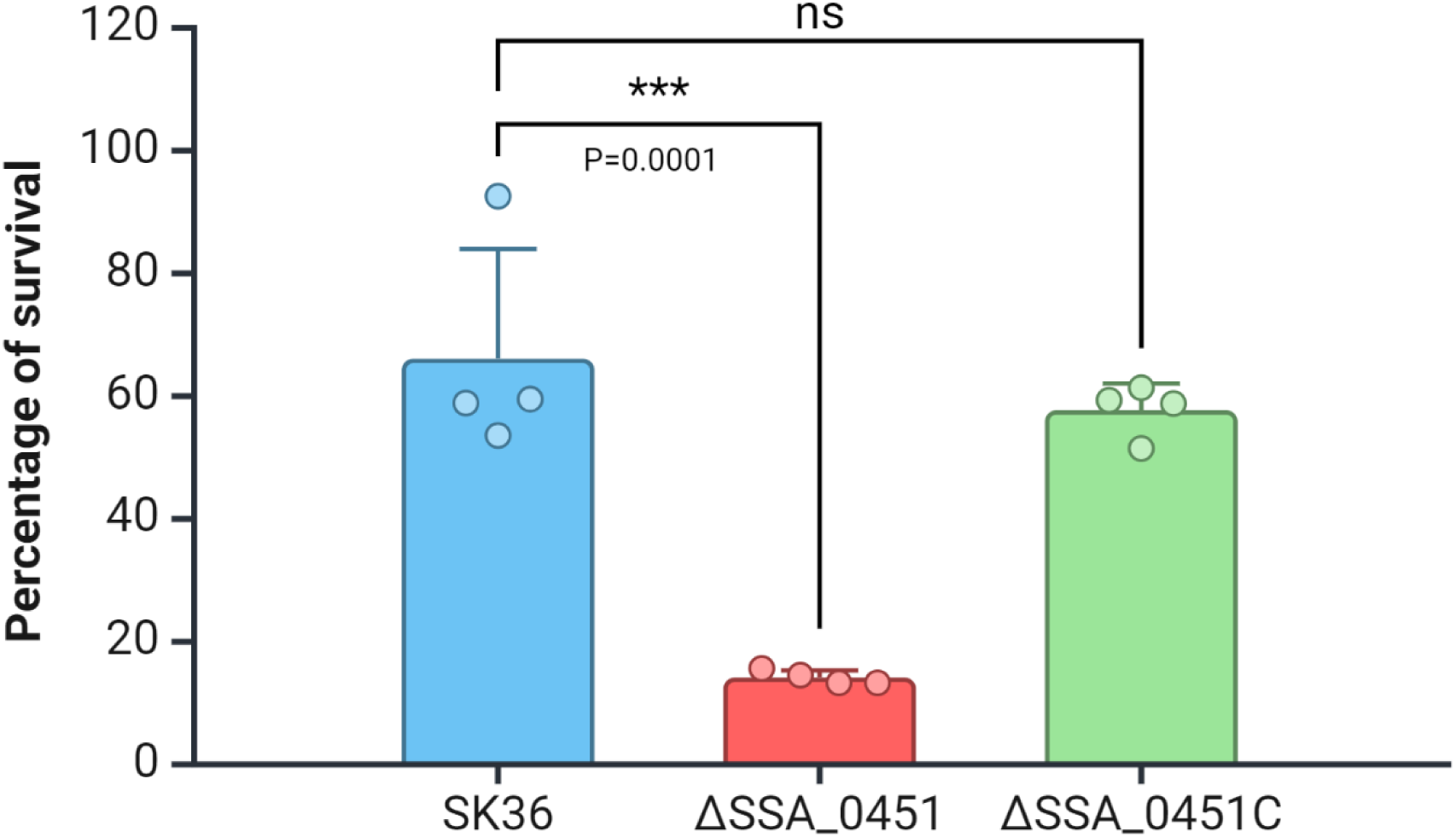
Survival of ΔSSA_0451 and ΔSSA_0451C after 30 min was shown based on CFU count. Values are ± SD, n=4.

**S1 Fig 3.**
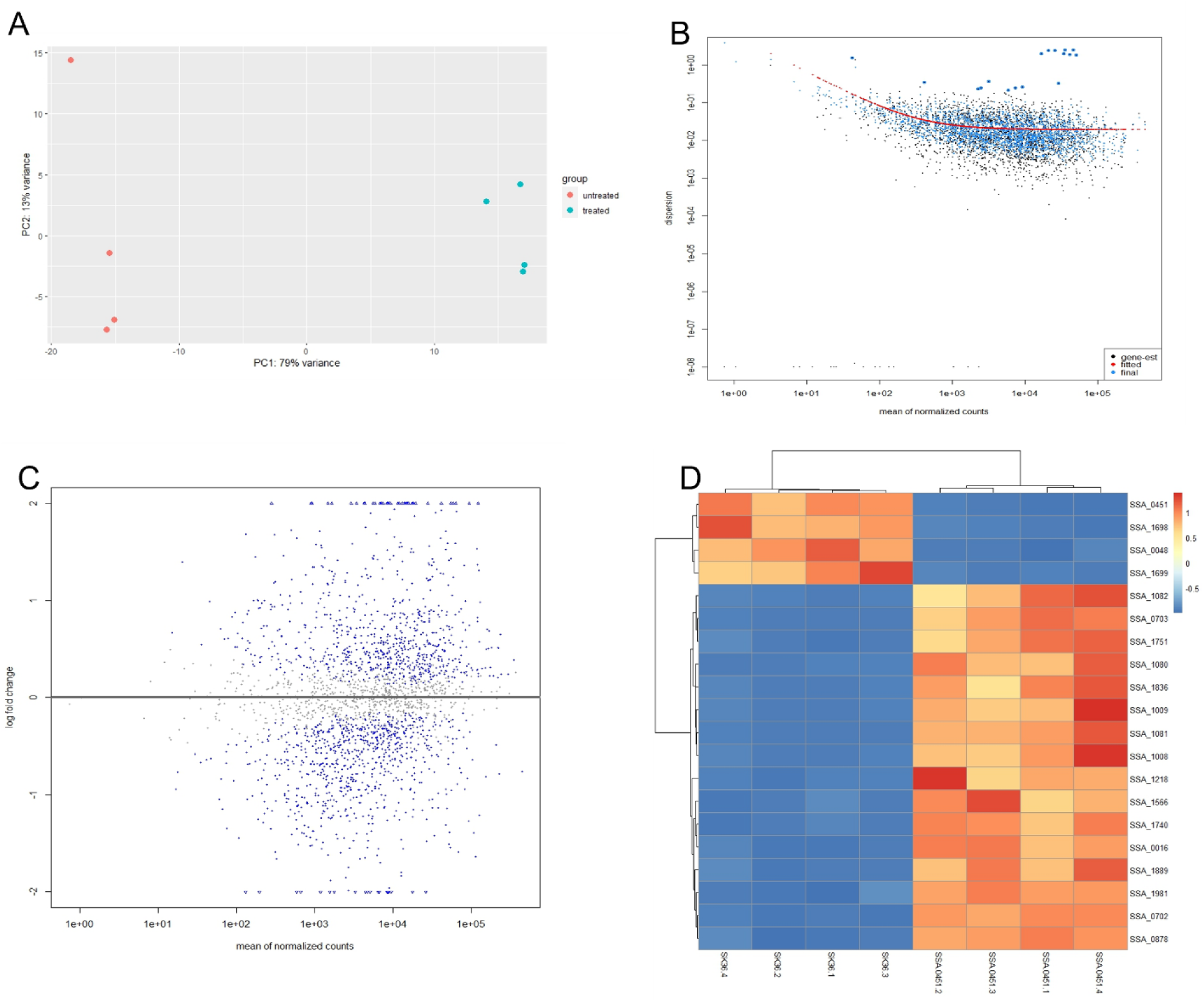
RNA-Seq analysis of ΔSSA_0451 and SK36 showed upregulated and downregulated genes. A: PCA plot shows the distribution of SK36 and ΔSSA_0451 biological replicates. B: Dispersion analysis of SK36 and ΔSSA_0451 based on mean normalized counts., C: MA Plot shows the distribution of upregulated and downregulated genes based on logfold change and mean normalized counts. D: Top upregulated and downregulated genes based on Z-score.

**S1 Fig 4.**
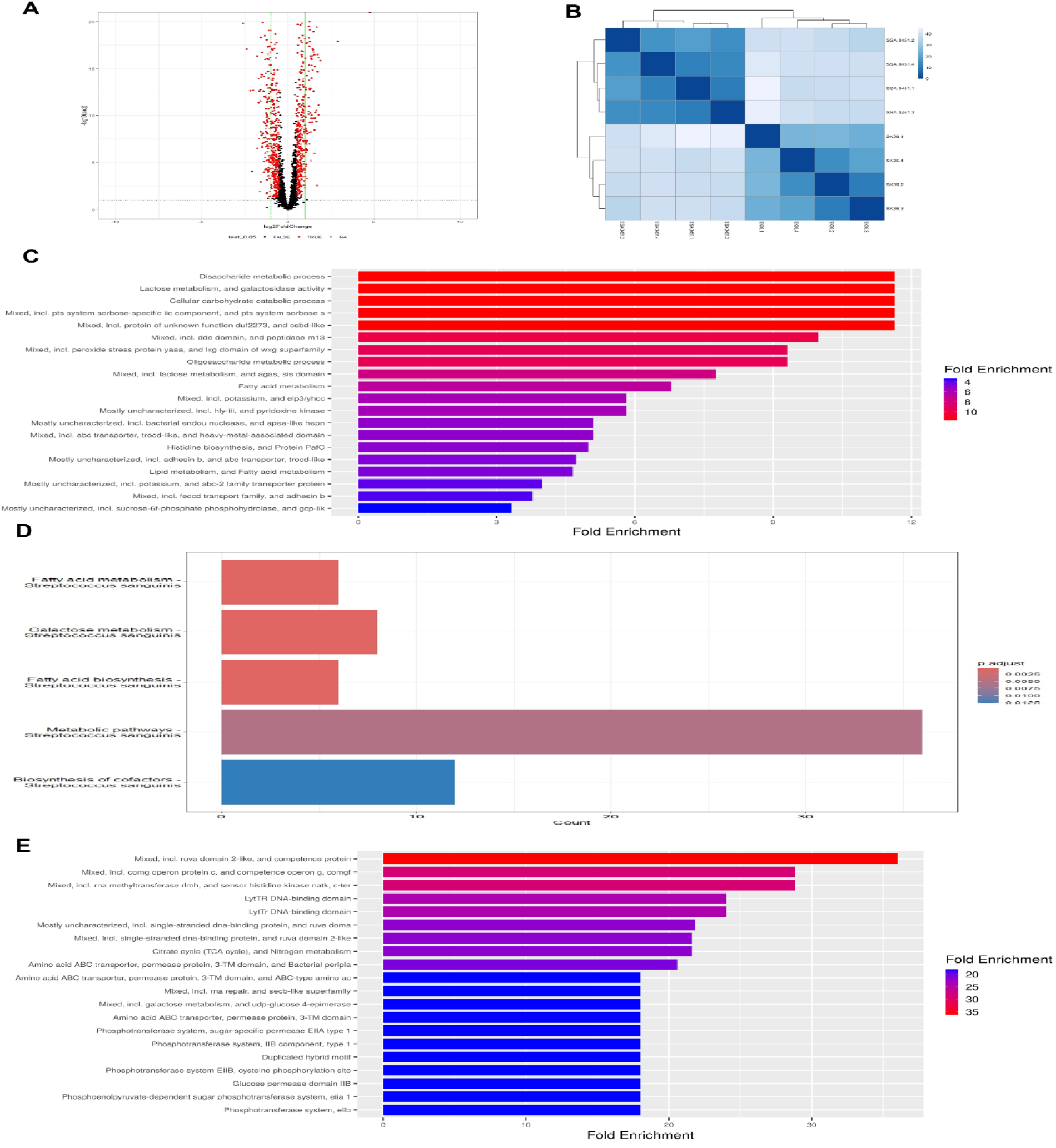
RNA-Seq analysis of ΔSSA_0451 and SK36. A: Volcano plot shows upregulated and downregulated genes based on their log2 fold value. B: Heat Map shows the difference between SK36 and ΔSSA_0451 replicates. C: The Gene ontology analysis of −1.5 log2fold genes shows the downregulated biological process. D: KEGG pathway analysis of −1.5 log2fold decreased genes shows the downregulated pathways. E: The Gene ontology analysis of 1.5 log2fold increased genes shows the upregulated biological process.

**S1 Fig 5.**
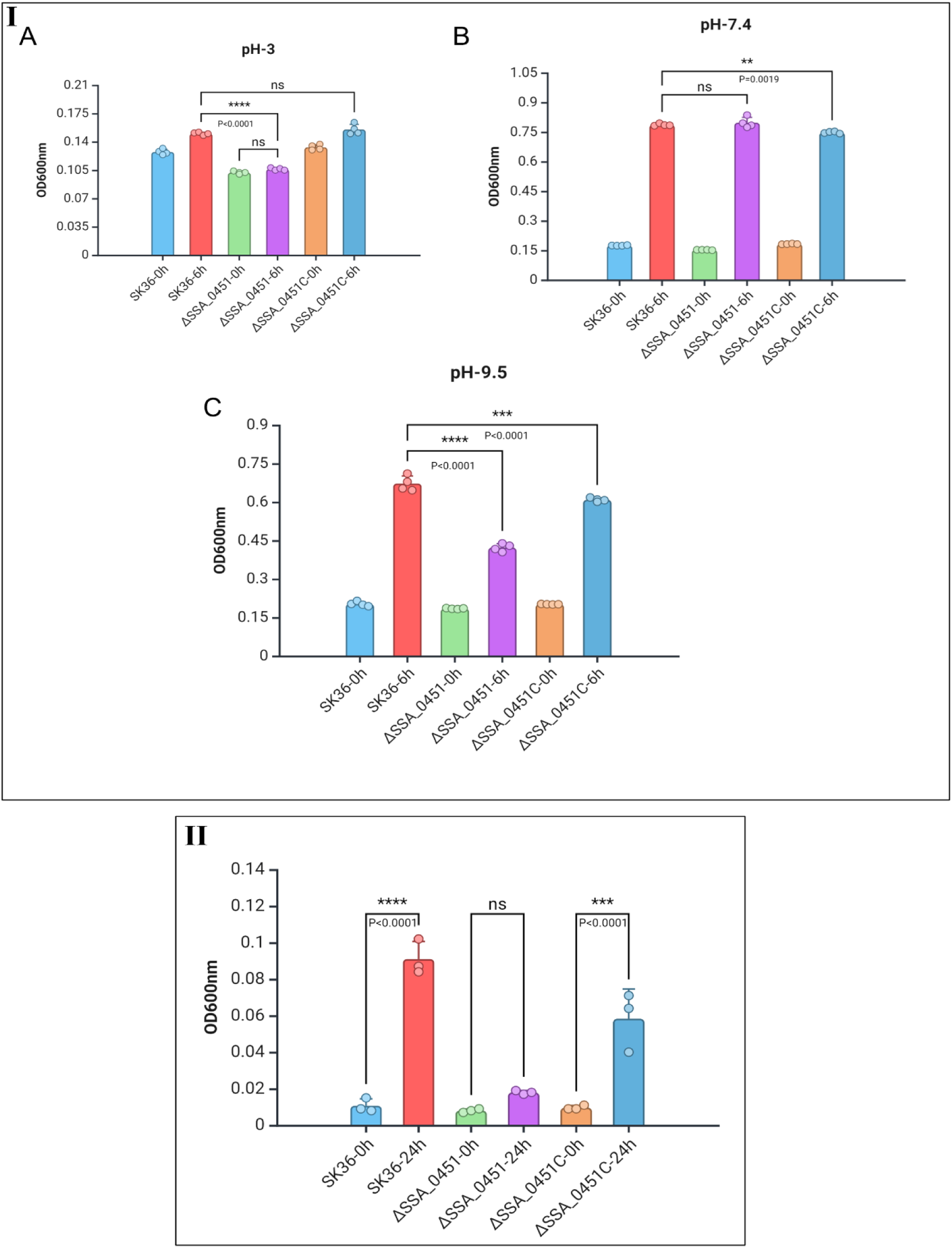
The SSA_0451 gene is necessary for *S. sanguinis* to survive under stress. I-A-C: Survival of ΔSSA_0451 in different pH. II: Survival of ΔSSA_0451 in bile salt.

**S1 Fig 6.**
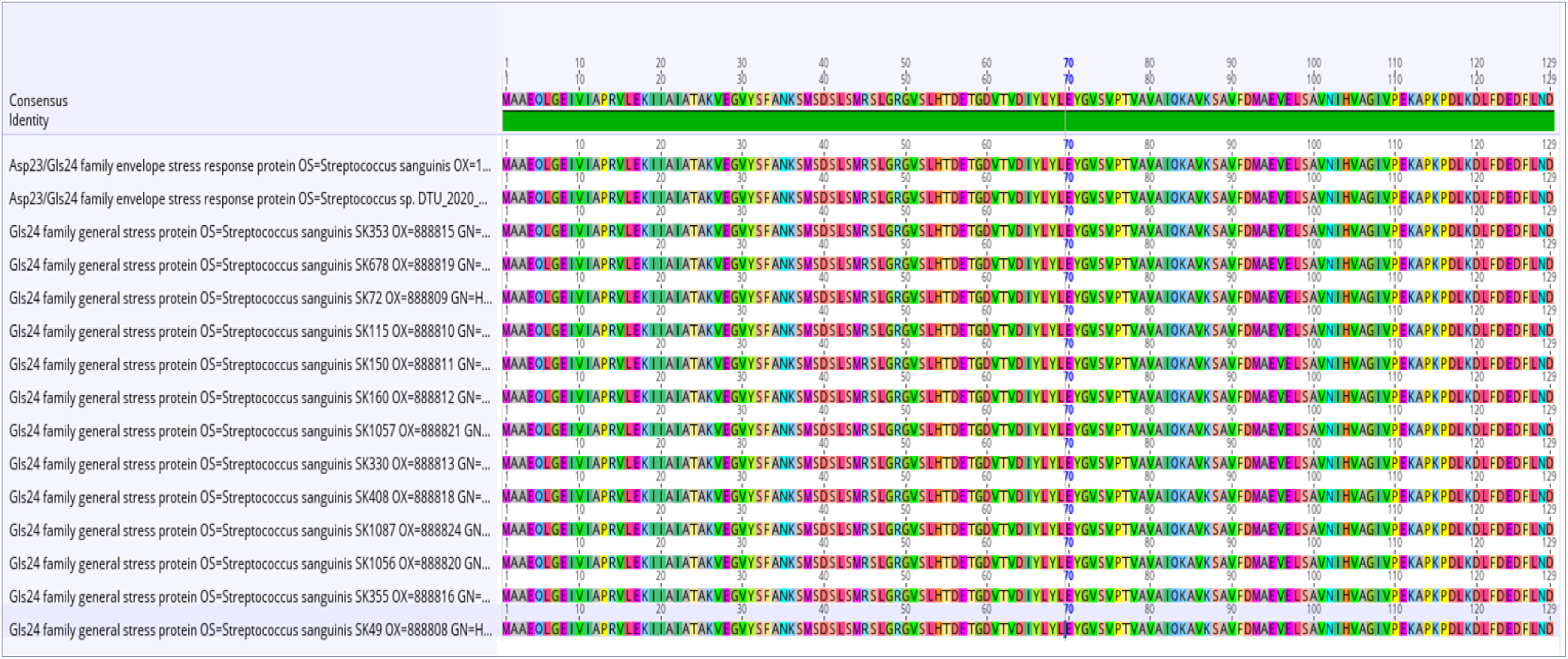
Protein homologous search. The Protein homologous search displayed that this protein is homologous to the alkaline shock protein-23 (Asp-23) or Gls24 family.

**S1 Table 1.** PTS system’s mannose-specific IIC component-related genes were downregulated in RNA-Seq analysis of ΔSSA_0451 and SK36.

| Locus | log2 foldChange | pvalue | padj |
| --- | --- | --- | --- |
| SSA_0219 | -3.20344 | 2.15E-83 | 2.10E-81 |
| SSA_0220 | -3.03231 | 8.50E-37 | 2.45E-35 |
| SSA_0221 | -2.86507 | 5.28E-36 | 1.43E-34 |
| SSA_0222 | -2.82302 | 5.85E-30 | 1.16E-28 |
| SSA_0224 | -2.92698 | 3.75E-22 | 4.87E-21 |
| SSA_2023 | -2.38917 | 8.11E-19 | 8.61E-18 |

The values were obtained from four biological replicates.

**S1 Table 2.** Strains and plasmids used in this study.

| Strain and plasmids | Description | Reference |
| --- | --- | --- |
| *S. sanguinis* SK36 | Human plaque isolate | [39] |
| JFP36 | Em<sup>r</sup>; Δ0169 : : pSerm, derived from SK36 | [49] |
| JFP56 | Sc<sup>r</sup>; Δ0169 : : aad9, derived from SK36 | [49] |
| ΔSSA_0451 | SK36, insert-deletion of SSA_0451 gene, Km<sup>r</sup> | [50] |
| pVMTeal-Plasmid | Gram-positive replicative plasmid carrying *S. mutans* ldh promoter and codon-optimized gene for mTFP1; Em<sup>r</sup> | [43] |
| pVMcherry-Plasmid | Gram-positive replicative plasmid carrying *S. mutans* ldh promoter and codon-optimized gene for mCherry; Em<sup>r</sup> | [43] |

Km, kanamycin; Em, erythromycin; Sc, spectinomycin.

**S1 Table 3.**
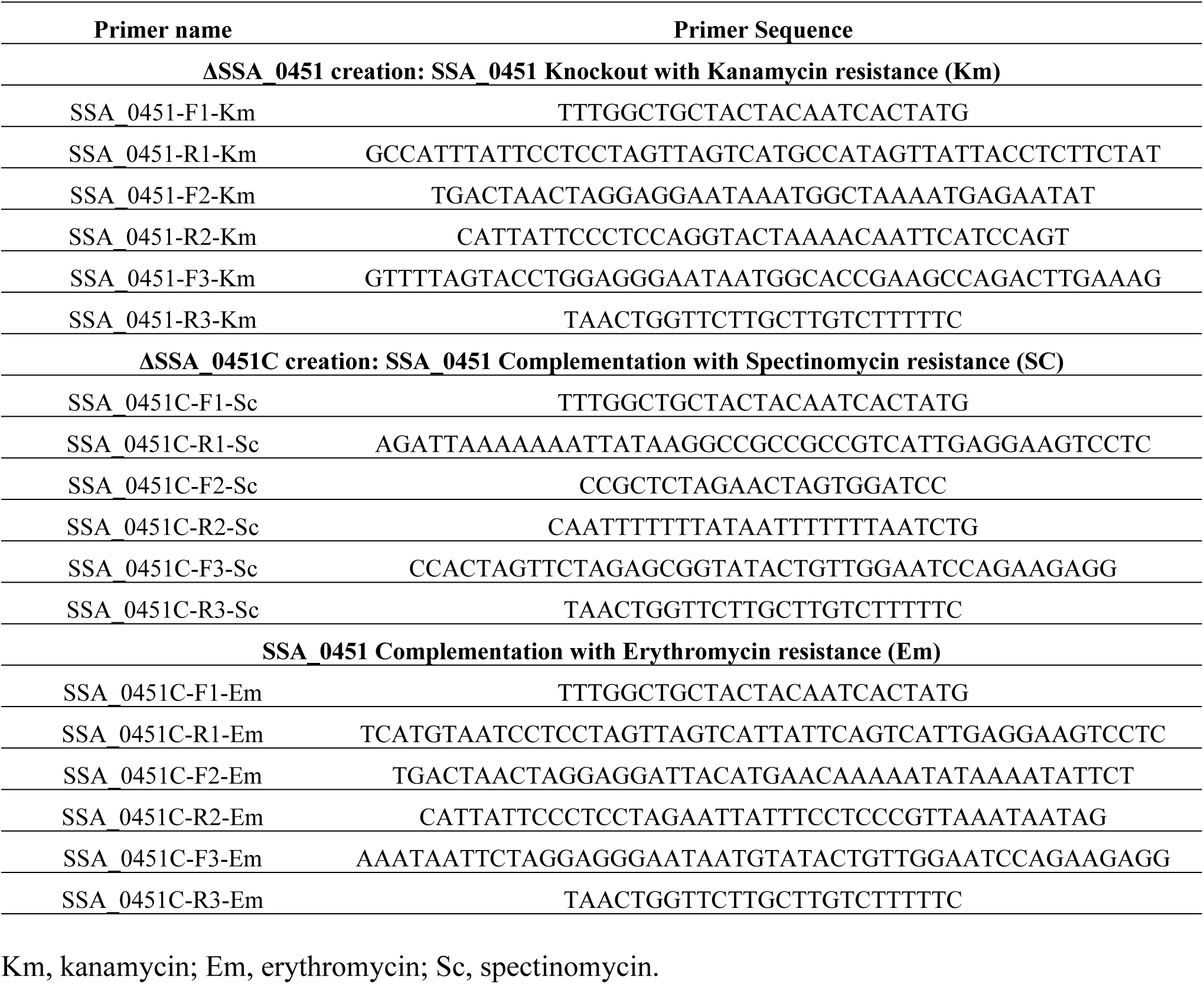
Primers used in this study.

**S1 Table 4.**
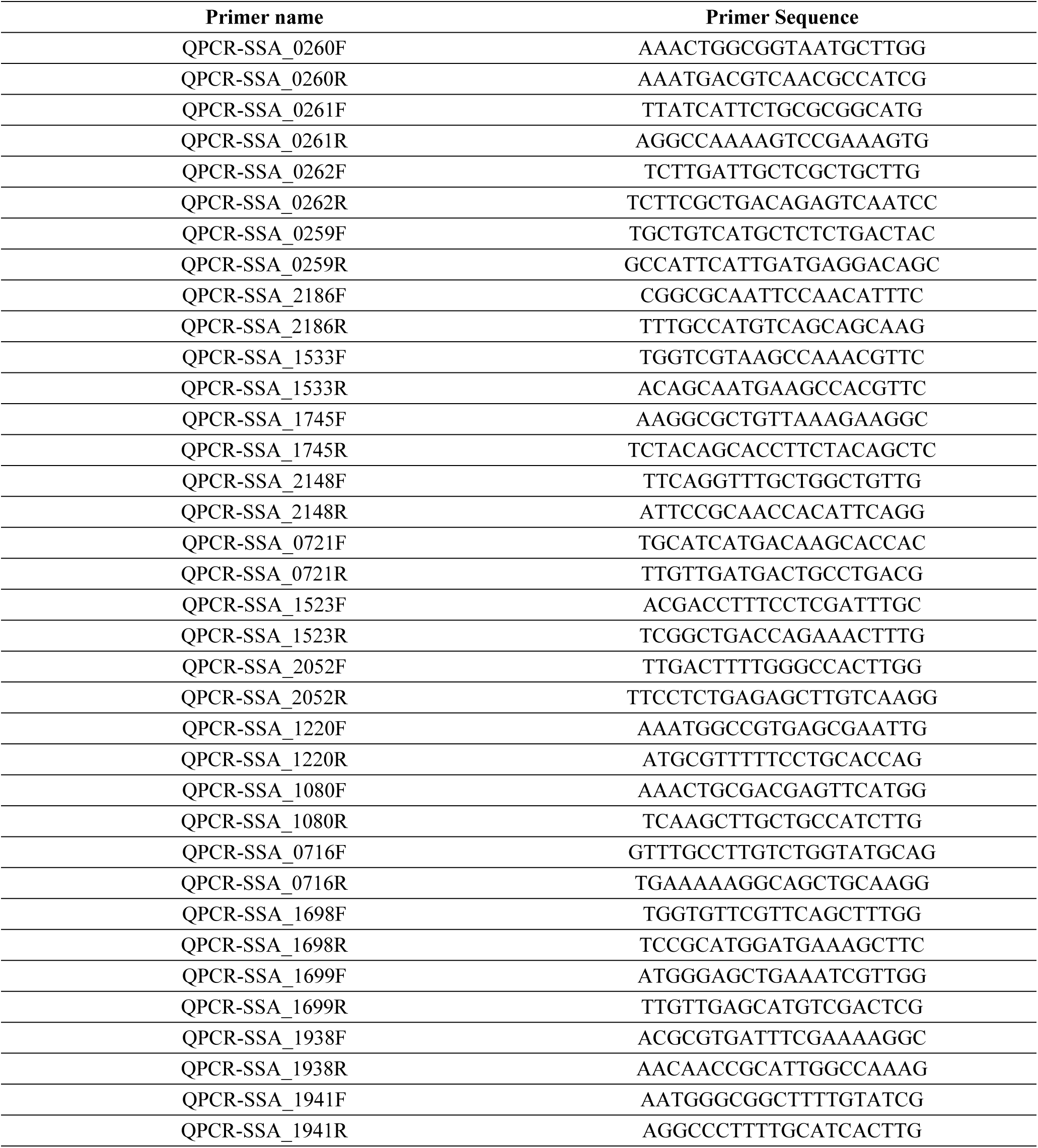
Primers used for qPCR study.

## Notes

### Competing Interest Statement

The authors have declared no competing interest.

